# The development of bi-directionally coupled self-organizing neurovascular networks captures orientation-selective neural and hemodynamic cortical responses

**DOI:** 10.1101/2021.12.24.474094

**Authors:** Bhadra S Kumar, Philip J. O’Herron, Prakash Kara, V Srinivasa Chakravarthy

**Affiliations:** Department of Biotechnology, Indian Institute of Technology Madras (IITM), India; Department of Physiology, Augusta University, Augusta GA; Department of Neuroscience, University of Minnesota, Minneapolis MN 55455, USA; Center for Complex Systems and Dynamics, Indian Institute of Technology Madras (IITM), India

## Abstract

The network of neurons in the brain is considered the primary substrate of information processing. Despite growing evidence on the possible role of cerebral blood flow in information processing, the cerebrovascular network is generally viewed as an irrigation system that ensures a timely supply of oxygen, glucose, and nutrients to the neural tissue. However, a recent study has shown that cerebral microvessels, like neurons, also exhibit tuned responses to sensory stimuli. Tuned neural responses to sensory stimuli are certainly enhanced with experience-dependent Hebbian plasticity and other forms of learning. Hence it is possible that the densely interconnected microvascular network might also be subject to some form of plasticity or competitive learning rules during early postnatal development such that its fine-scale structure becomes optimized for metabolic delivery to a given neural micro-architecture. To explore the possibility of adaptive lateral interactions and tuned responses in cerebral microvessels, we modeled the cortical neurovascular network by interconnecting two laterally connected self-organizing networks (Laterally Interconnected Synergetically Self-Organizing Map - LISSOM). The afferent and lateral connections of the LISSOM were defined by trainable weights. By varying the topology of lateral connectivity in the vascular network layer, we observed that the partial correspondence of feature selectivity between neural and hemodynamic responses could be explained by lateral coupling across local blood vessels such that the central domain receives an excitatory drive of more blood flow and a more distal surrounding region where blood flow is reduced. Critically, our simulations suggest a new role for feedback from the vascular to the neural network because the radius of vascular perfusion seems to determine whether the cortical neural map develops into a clustered and columnar vs. salt-and-pepper organization.

## Introduction

It has been long-known that neurons that exhibit tuned responses to various visual properties like orientation, direction, ocular dominance, color[4–6], and complex form[7,8] are present in various visual cortical areas in the brain. Responses to oriented stimuli have been observed even in the astrocytes of the visual cortex[9]. Taking a step further, O’Herron and colleagues [10] observed that even the microvessels in V1 exhibit tuned responses to oriented moving gratings, even though the specificity of the response to the preferred stimulus is not as sharp as that of the neurons. More specifically, in a given orientation column of neurons, even though an individual arteriole responded maximally to the same preferred orientation as all of its neighboring neurons, that blood vessel also responded to other stimulus orientations, whose neural signals had to come from adjacent orientation columns.

Computational studies have shown that a neural network is capable of generating tuned responses to input stimuli by means of competitive learning, which can be implemented in a neural network by lateral interactions among the neurons characterized by long range inhibitory signaling and local excitatory signaling, often described as an ON-Center, OFF-surround neighborhood interaction[11–13]. The existence of the lateral excitatory and inhibitory connections among cortical neurons was supported by experimental studies[14–17]. A recent study shows the relevance of tightly coupled excitatory and inhibitory networks in the developing visual cortex with self-organizing ability. These various experimental findings led us to determine if competitive learning mechanisms could also underlie the generation of tuned responses in cerebral microvessels.

O’Herron et al. [10] presented data which suggested that the reduced selectivity in the vessels (as compared to the neurons in their perfusion field) was likely due to long-range propagation of dilation from one cortical column to a neighboring column. But other mechanisms, none of which were tested, are also possible. For example, the long-range release of vasodilator, or the modulation of arteriolar diameter by transmitters released from inhibitory neurons and the release of vasodilatory substances by astrocytes, are some plausible mechanisms. We argue that the effect brought about by all the aforementioned biological mechanisms of vessel-to-vessel interactions can be computationally visualized as manifestations of lateral interactions among the vessels. The arterioles with their ability to release nitric oxide, a well-known vasodilator[18], can directly mediate lateral interactions among the vessels. Propagation of vasodilatory signals along endothelial gap junctions is yet another source of long-range signaling among the vessels[19–22]. Likewise, the ability of interneurons and astrocytes to mediate vasodilation and vasoconstriction [23–26] can also be invoked as an indirect basis for lateral interactions among the vessels.

Tuned responses in vessels could suggest the existence of some kind of competitive learning mechanism in the vascular network. Computational models that implement ON-Center, OFF-surround neighborhood interactions have been successful in describing topographic maps in the brain, not just in the visual cortex, but in several other sensory cortical areas as well [19, 21]. It would be worthwhile to investigate if ON-Center, OFF-surround neighborhood interactions among microvessels of the visual system can result in tuned responses. We therefore explored these possible mechanisms by using a computational model of the neurovascular network in the visual cortex.

To investigate the presence of competitive learning in the cerebrovascular layer, we propose a computational model of the development of a neurovascular network where both neural and vascular layers are described as self-organizing networks, each with its own lateral connectivity. The model is termed ‘Neuro Vascular coupling using Laterally Interconnected Networks’ (NV-LIN) (fig.1). NV-LIN explores the different lateral connectivity patterns in the vascular layer to identify the best configuration of the lateral interactions in the vascular layer that produces tuned responses in the single vessel that match the tuned vascular responses in the experimental study [10]. To simulate the neural network in V1 of the non-rodent cortex with a columnar organization for orientation selectivity, we used a biologically plausible self-organizing network architecture like the Laterally Interconnected Synergetically Self-Organizing Map (LISSOM) architecture [27].

**Figure 1:**
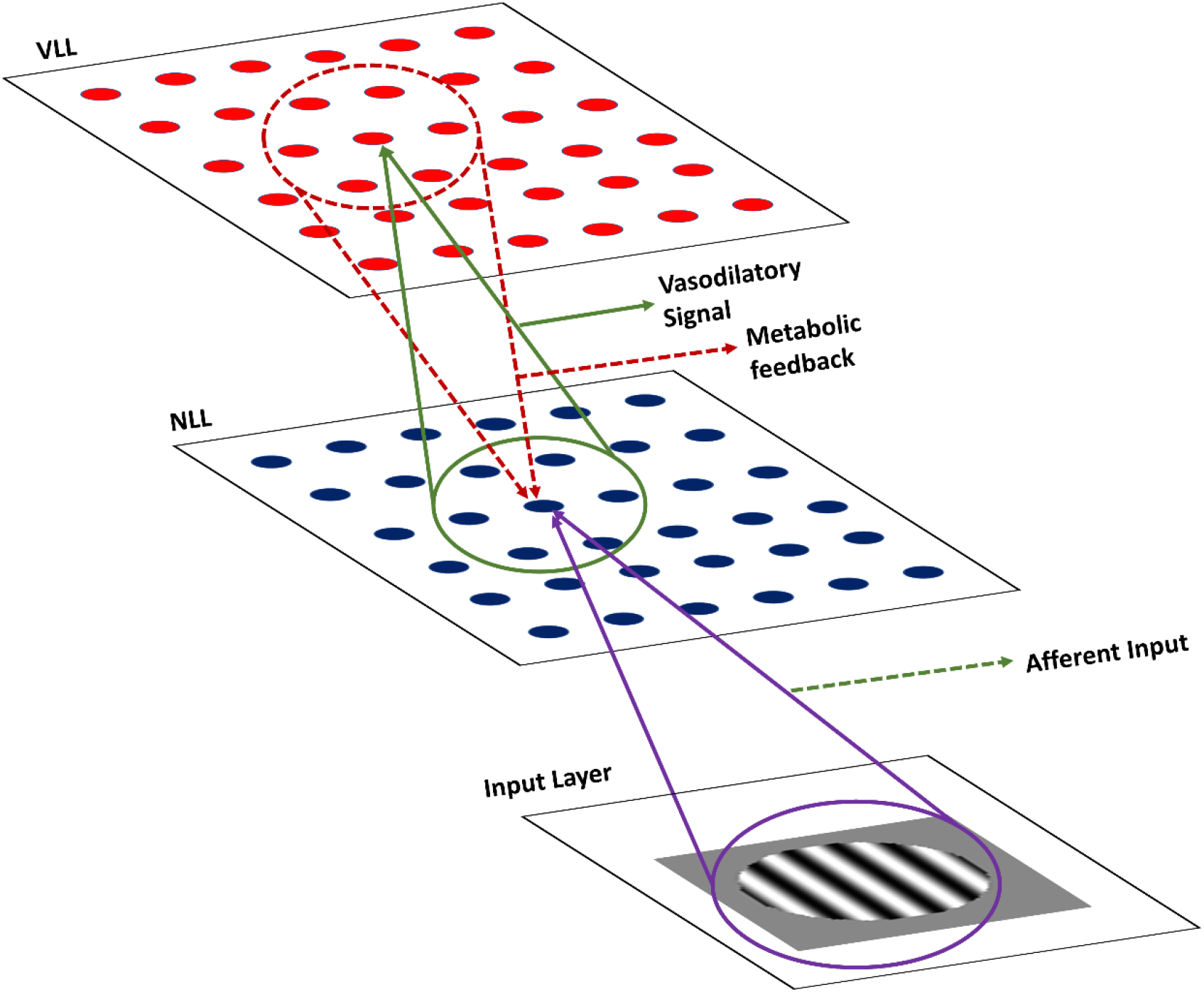
A Schematic Representation of the architecture of the NV-LIN Model. Neural LISSOM Layer (NLL) is bidirectionally connected with Vascular LISSOM Layer (VLL). Each unit of NLL receives the weighted sum of a region of the moving grating stimulus from its receptive field in the input layer and also receives metabolic feedback from the vascular units in VLL, which perfuse them. Each unit of VLL receives the vasodilatory signal from the neural units in its receptive field in the NLL.

An assumption that forms the basis for the interpretation of functional neuroimaging techniques [26,28,29] is that functional hyperemia observed in cerebral blood vessels is largely a consequence of the proximal neural activity[28,29]. In our recent work, where a rate-coded neural network (modeled using LISSOM) was bidirectionally coupled with a vascular anatomical network model, it was observed that the spatial distribution of hemodynamic response pattern closely follows the proximal neural firing rate (see fig.6A and B in Ref. [30]). Since our current goal is to account only for the spatial distribution pattern of the hemodynamic response, we propose modeling the vascular response using a network with fundamental units similar to that of LISSOM (sigmoid neural units). This approach simplifies the computational complexity required to design a bidirectionally coupled model, where the lateral connectivity architecture of one of the layers (vascular layer in this case) needs to be explored by trying all possible topologies.

A LISSOM used to simulate the neural layer is well known to have a lateral connectivity pattern with near excitatory (ON center) and far inhibitory (OFF surround) connections[27,31]. But this is not the case when the network is used to simulate the vascular layer. Since the lateral connectivity pattern among the blood vessels is unknown, we explore several possible topologies including, for example, OFF-center ON-surround, to identify the lateral connectivity pattern which would closely approximate the experimentally observed vascular responses.

Our new NV_LIN model architecture consists of 3 layers: input layer, neural layer and vascular layer (see fig. 1 and Methods). The input layer represents the visual input that is presented to the neural layer. The neural layer, in turn, projects to the vascular layer. In biological terms, these feedforward connections could be treated as lumped representations of the many ways in which neurons activate proximal microvessels[32]. For example, directly by the release of nitric oxide (NO)[33–35] from neurons, the release of a neuromodulators like dopamine[36] and indirectly by activating astrocytes [37,38] which release arachidonic acid (AA) metabolites like EET and PGE2 [39–42]. The neurovascular coupling in our NV_LIN model was designed by also keeping in mind the proof-of-concept ideas proposed in the hemoneural hypothesis[1]. Many experimental observations[2,3], as well as the results from our previous computational models [30,43,44] support the hypothesis by Moore and Cao[1] that hemodynamics could impact neural activity directly or indirectly. Hence in our NV_LIN model, the vascular layer is designed to send feedback to the neural layer. These feedback connections can be thought to represent one of many ways in which vessels can influence neural activity – release of metabolic substrates like glucose and oxygen that directly sustain neural activity[45], the release of the same metabolic substrates to astrocytes, thereby indirectly influencing neural activity[46–48], mechano-sensation of changes in arterial wall diameter, a mechanism that has been dubbed vasculo-neural coupling [2]. In view of this changing perspective of neurovascular coupling, we introduce bidirectional coupling between neural and vascular layers in NV_LIN model.

In the first phase of our study, our NV-LIN model was simulated under three different lateral connectivity topologies in the vascular layer: (i) No lateral connectivity, (ii) ON center OFF surround and (iii) OFF center ON surround. The bidirectional connectivity between the neural and vascular layers in NV-LIN simulates the bidirectional neurovascular coupling. The vascular response depends on the afferent neural input, and this neural input, in turn, depends on the feedback from the vascular network, in addition to the external stimulus. The tuned response of neural and vascular units was compared with the experimental data to find the optimal lateral connectivity topology in the vascular network.

In the second phase of our study we examined whether vascular feedback influences the specific kind of neural map architecture that may be formed. This is because of the known effects of the vasculature in maintaining neural firing rates (see earlier discussion paragraphs). More broadly, the functional significance of clustered neural maps in the cerebral cortex is poorly understood, where the default role is largely ascribed as an epiphenomenon of development in minimizing neuronal wiring. We addressed the role of vascular feedback on neural map formation in our model by altering the vascular feedback to the neural network in terms of the perfusion field of the vessel while keeping the sensory input to the neural layer and the network parameters like excitatory radius, inhibitory radius, learning rate and scaling parameters unchanged. We show that the type of neural functional architecture that develops, can be dramatically influenced by a change in vessel perfusion field.

## Methods

The model consists of two layers of a bidirectionally connected LISSOM network (fig. 1). The LISSOM network is a biologically plausible self-organizing network capable of generating tuned responses to input stimuli at individual unit level and organizational maps of the input stimuli at a global level. The first LISSOM layer represents the neural layer and is named the Neural LISSOM Layer (NLL), and the second LISSOM network represents the vascular layer and is named the Vascular LISSOM Layer (VLL). The network was trained using moving sinusoidal grating stimuli of different orientations and a temporal frequency of 2 Hz. The gratings were presented in eight different directions in increments of 45°. A total area of 3mm x 3mm from the V1 area in the visual cortex was considered. The receptive field of the vessels, which indicates how far a neuron can cause vasodilation in that vessel, was fixed to be close to 430μm. We fixed this value by assuming that the vasodilatory signals resulting from neural activity could impact vessels in approximately same area equal to the typical perfusion field of a vessel (~400μm[49]). The perfusion area of a vessel was fixed to be ~430 μm in the model. The VLL was provided with a constant input to simulate the baseline vascular tone. The network was evaluated by considering the activity of each vascular unit and the average neural activity of the neural units in the perfusion domain of that vessel (~430 μm).

In the neural layer, each unit receives afferent connections from the input, lateral connections from the neighboring neurons, and feedback connections from the vascular layer (fig 2. a). The total input 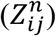 received by the neural unit at location (*i,j*) is given by

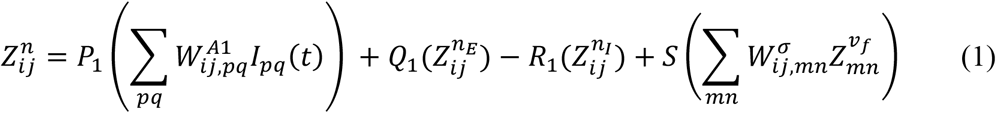

where P_1_, Q_1_, R_1_, and S are constants, *W*^*A*1^ is the afferent weight stage from the input layer I to the neural layer. 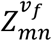 is the activity at the vascular node (*m, n*) and the vasculo-neuronal feedback weights, *W^σ^* is denoted by a two-dimensional Gaussian function representing the diffusion of the energy substrates from the vessels to the neural tissues. The width of the isotropic Gaussian curve depends on the perfusion field defined for the vessel. The perfusion field of a vessel indicates the field over which a vessel can perfuse neurons in the neural layer. 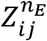 and 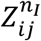 are the lateral inputs from the neighboring excitatory neural units and the inhibitory neural units, respectively, and are estimated as follows.

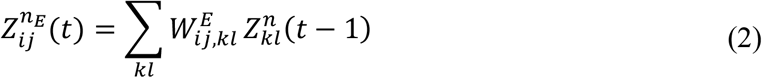

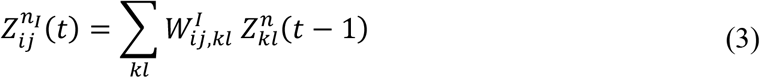

*W^E^* are the weights connecting the excitatory neighborhood and *W^I^* are the weights connecting the inhibitory neighborhood. (*i,j*) denotes the index of a neural unit in the two-dimensional grid; (*k, I*) denotes the index of a neural unit in the neighborhood, which gives excitatory or inhibitory projections to the neural unit at location (*i,j*) depending on their proximity.

**Figure 2:**
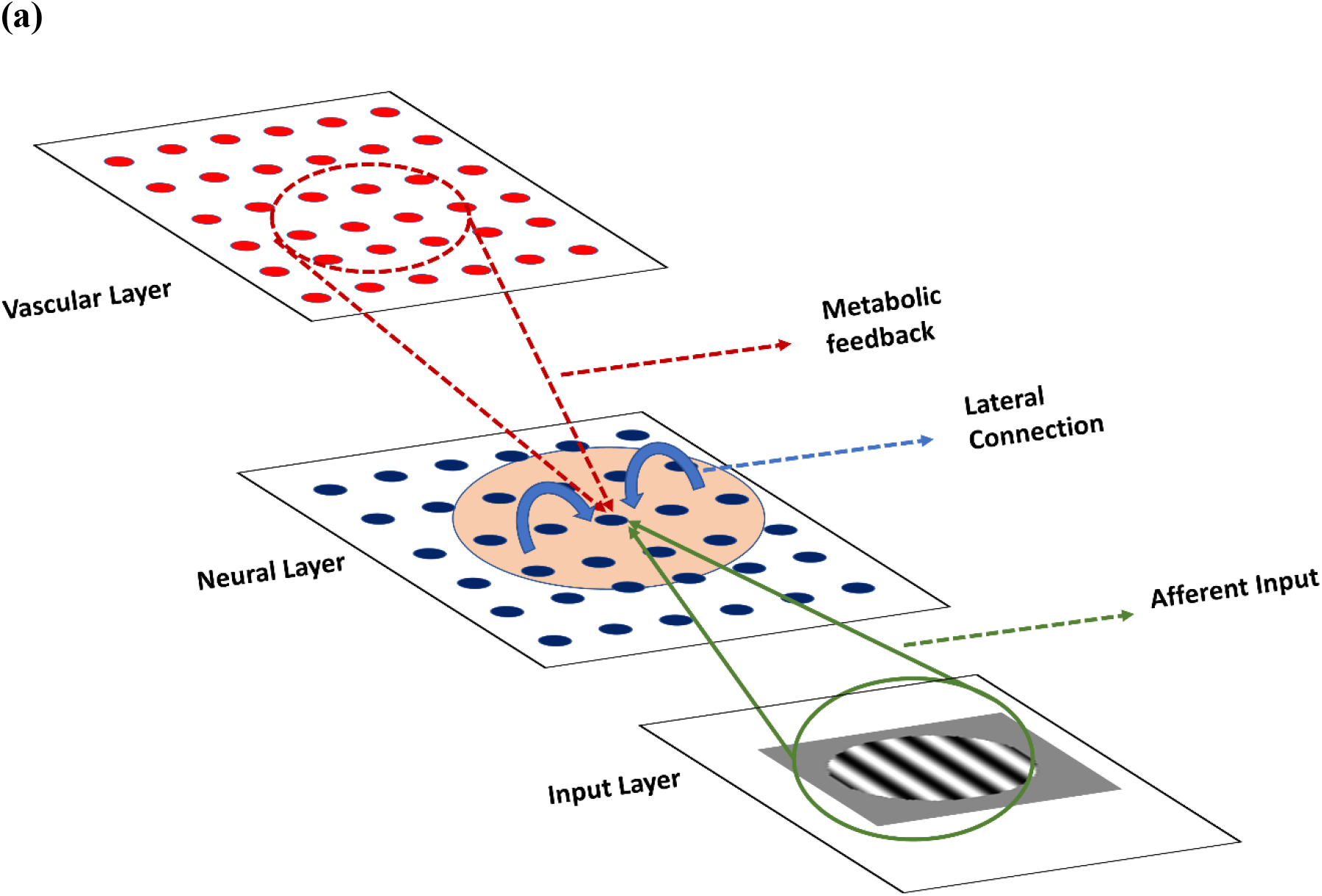
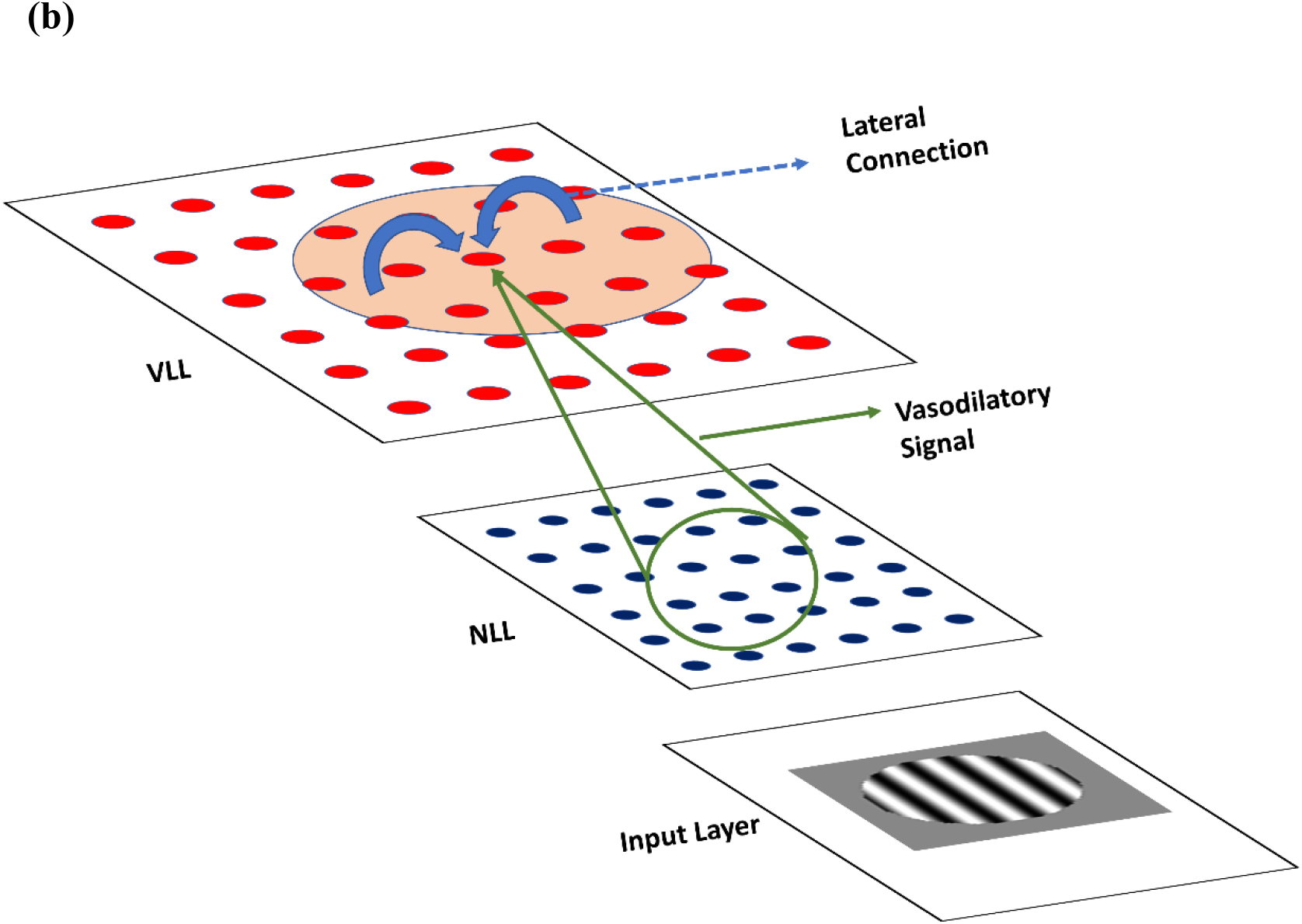
(a) Inputs received by a unit in the NLL (b) Inputs received by a unit in VLL.

Similarly, the total input 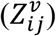 received by the vascular unit at location (*i,j*) is given by (fig.2b)

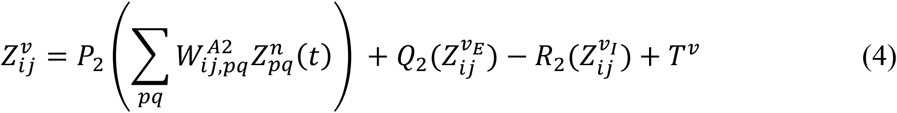

where *P*_2_, *Q*_2_, *R*_2_, are constants, *W*^*A*2^ is the afferent weight stage from the NLL to the VLL and 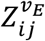 and 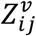 are the inputs from the neighboring excitatory vascular units and the inhibitory vascular units, respectively. *T^v^* is a constant representing the minimum constant tone of the vessels. The receptive field of each vessel was defined in such a way that a neuron as far as 600 μm from it could send a vasodilatory signal to the vessel.

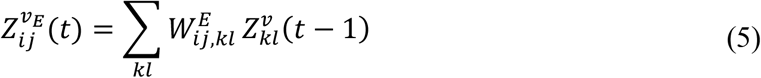

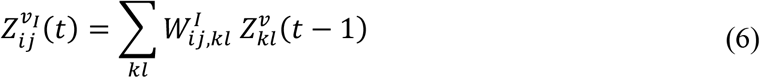

*W^E^* are the weights connecting the vessel to its excitatory neighborhood and *W^I^* are the weights connecting the vessel to its inhibitory neighborhood. (*i,j*) denotes the index of a vascular unit in the two-dimensional grid; (*k, I*) denotes the index of a vascular unit in the neighborhood, which has excitatory or inhibitory connections with the vascular unit at location (*i,j*) depending on their proximity.

The weighted sums of inputs (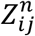 and 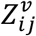) are passed through an activation function, in this case, a piecewise-linear sigmoid function (*σ*),

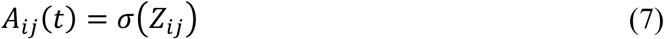

where *σ* is defined as,

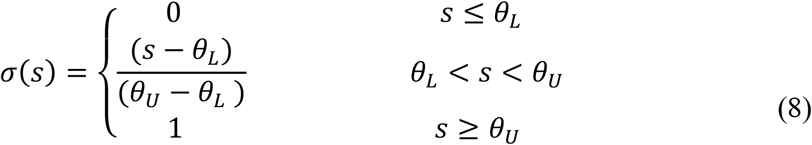

and the parameters *θ_L_* and *θ_U_* represent the transition points between the three branches of the sigmoid function.

The activity of the LISSOM layer is allowed to settle for each frame of the input. The afferent and lateral weights of both NLL and VLL are trainable. These weights are updated after every frame. The afferent weights to both NLL and VLL are trained using symmetric Hebbian learning, and the lateral connections are trained using asymmetric Hebbian learning. Given that an afferent weight 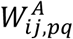 connects node (*i,j*) to node (*p, q*) and that Z is the activity of a given node, the weight is updated using symmetric Hebbian learning defined as,

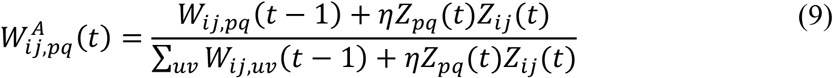

Given that a lateral weight 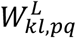 connects node (*k, l*) and node (*p, q*) and that Z is the activity of a given node, the weight is updated using asymmetric Hebbian learning defined as;

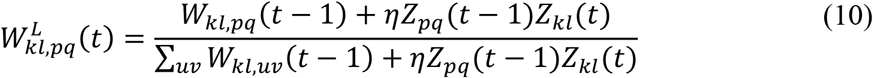

Note that whereas in the case of symmetric Hebbian learning, the pre- and post-activations are considered at the same time instant, in the case of asymmetric Hebbian learning, the pre-activation is considered at the previous time instant (t-1), and the post-activation is considered at the current (t) time instant.

The model performance was evaluated using two metrics, (i) Directionality Index (DI) and (ii) Orientation Selectivity Index (OSI). Directionality Index (DI) quantifies the ability of a unit to differentiate the direction of the motion among similar orientations,

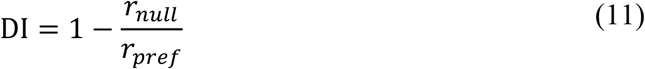

where *r_null_* is the response of the unit to the stimulus with the same orientation drifting in the opposite direction and *r_pref_* is the response of the unit to the preferred stimulus.

Orientation Selectivity Index (OSI) measures how specific the response of a neuron or a vessel is to a given orientated stimulus.

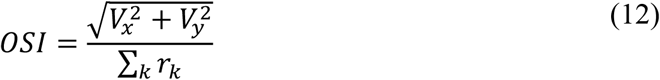

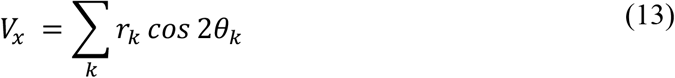

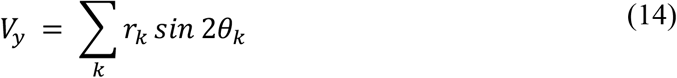

where *θ_k_* is the orientation of the *k*’th stimulus and *r_k_* is the response of each unit to that stimulus.

The Orientation Preference (OP) identifies the orientation to which the neuron or the vessel responds maximally and is calculated as,

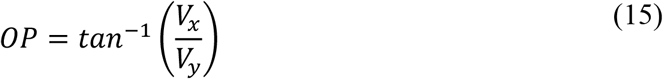

where *θ_k_* is the Orientation of the *k*’th stimulus and *r_k_* is the response of each unit to that stimulus.

The direction sensitivity of the neurons and vessels are indicated by means of a direction map that labels each unit in the LISSOM as the direction to which it responds maximally. To understand the nature of map formation in the neural network in response to change in the vascular perfusion field, we defined two metrics to evaluate the maps. (i) Number of Active Units (NAU) and (ii) Mean Area of Active Units (MAAU). For higher values of NAU, the response map tends to have a salt and pepper in nature.

The settled response of NLL is passed through a hard thresholding filter to obtain a binary image. NAU is then estimated by running connected component analysis [50,51] on these settled binary images. The number of connected components gives the value of NAU. The mean area of the NAU determines the MAAU.

**Table 1:** Values and units of constants.

| **Table 1: Values and units of constants** |  |
| --- | --- |
| **Constants** | **Value** |
| Size of neural layer | 63x63 |
| Size of vascular layer | 27x27 |
| Receptive field from input layer to neural layer | 9x9 |
| Receptive field from neural layer to vascular layer | 9x9 |
| Stride in neural layer | 1 |
| Stride in vascular layer | 2 |
| [P1, Q1, R1, S] | [0.4,1,1,0.6] |
| [P2, Q2, R2] | [0.5,0.8,0.3] |
| Excitatory Radius in neural layer | 5x5 |
| Excitatory Radius in vascular layer | 6x6 |
| Inhibitory Radius in neural layer | 15x15 |
| Inhibitory Radius in vascular layer | 15x15 |
| Radius of perfusion field | 9x9 |
| Threshold for connected component analysis | 0.2 for neural layer |

## Results

The network was studied in two phases. In the first phase, the possibility of lateral connectivity in vessels was investigated. The experimentally observed values of OSI and DI in neural and vascular networks were used to validate the model. In the later part of the study, the model was explored to predict the influence of the vascular perfusion field on the neural network map formation.

### The lateral connectivity architecture in the vessels

In computational neural models, orientation selectivity of neurons often arises from the competitive learning facilitated by the distant lateral inhibitory connections[27]. Latest experimental studies also support the existence of such long range inhibitory connections among cortical neurons facilitating the self-organizing ability of the cortex during the developmental stage[16]. Since the vascular network also exhibits tuned responses, it indicates the possibility of inhibitory lateral connections among the vessels, which enable competitive learning.

Three different lateral connectivity patterns were assigned to the VLL in order to identify the ideal pattern: (i) No lateral connectivity, (ii) ON-center, OFF-surround (iii) OFF-center, ON-surround. In the first case of “No lateral connectivity,” each vessel receives only afferent signals from the neurons and a constant input (fig. 3.a). In the second case of “ON-center, OFF surround,” in addition to the afferent signals from neurons, a vessel receives excitatory signals from its immediate neighborhood, and receives inhibitory signals from the vessels that reside in the annular region beyond the excitatory region (fig 3.b). In the third case of “OFF-center, ON-surround,” in addition to the afferent signals from neurons, a vessel receives inhibitory signals from its immediate neighboring vessels, and excitatory signals from the vessels located in the annular region beyond the inhibitory region (fig 3.c).

**Fig. 3.**
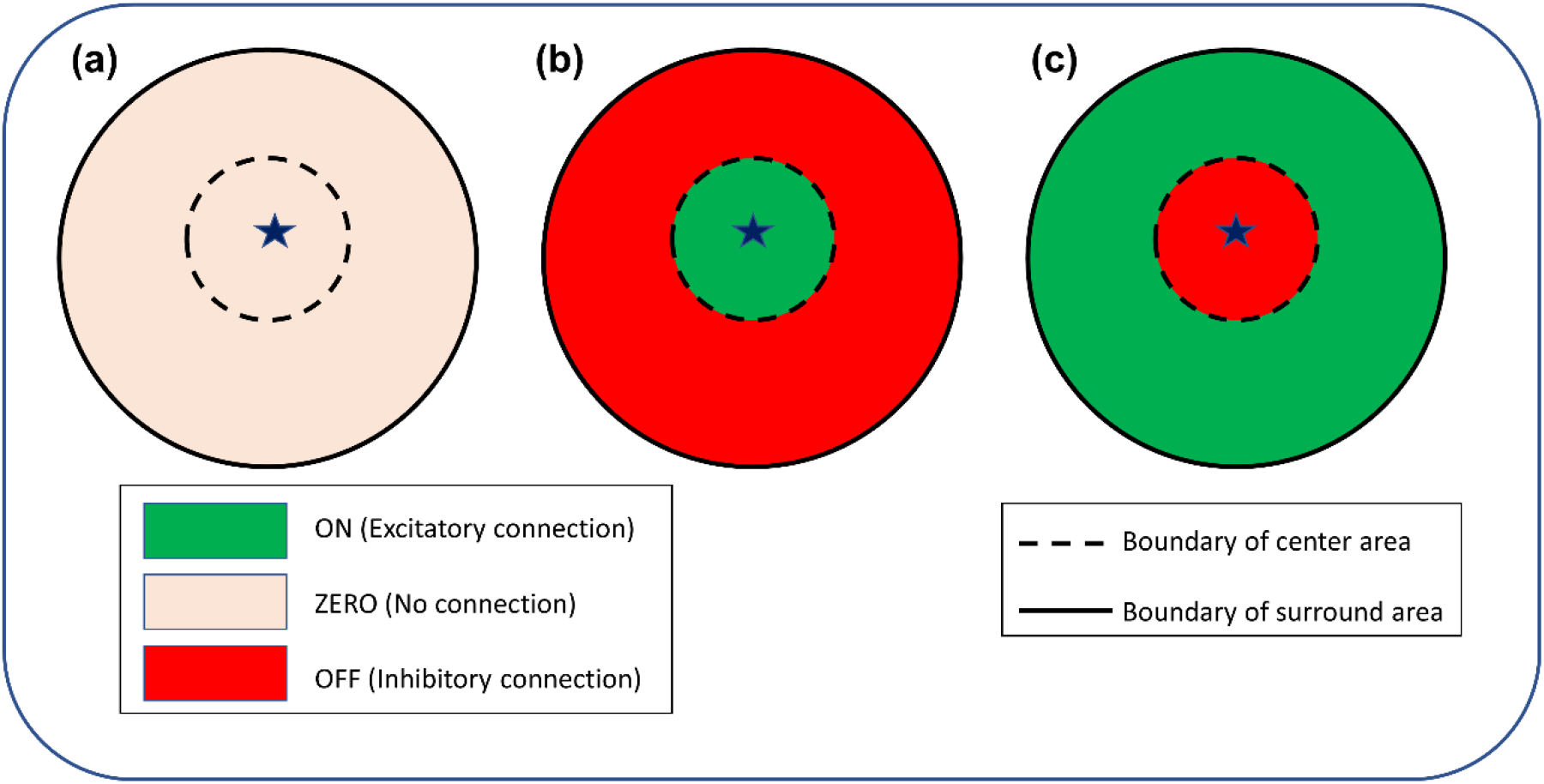
Schematic of the possible lateral connectivity architectures in the VLL. (a) No laterals (b) ON-center, OFF-surround (c) OFF-center, ON-surround.

The tuned responses of the NLL and VLL to oriented moving gratings were evaluated in terms of OSI and DI. Each unit in the VLL was given a lateral connection with a center-surround topology. For a fixed central region of 6×6 units and a surround region of 15×15 units, the lateral connectivity pattern was varied over the three defined conditions (Fig 3). The architecture was varied by varying the signs of the scaling factors Q2 and R2 in equation (4) accordingly.

The OSI and DI exhibited by all the three vascular architectures were compared with the experimentally observed tuned response indices, as shown in fig. 4. The VLL with ON center OFF surround later connectivity architecture gave the evaluation metric closest to experimental observations[10].

**Fig. 4:**
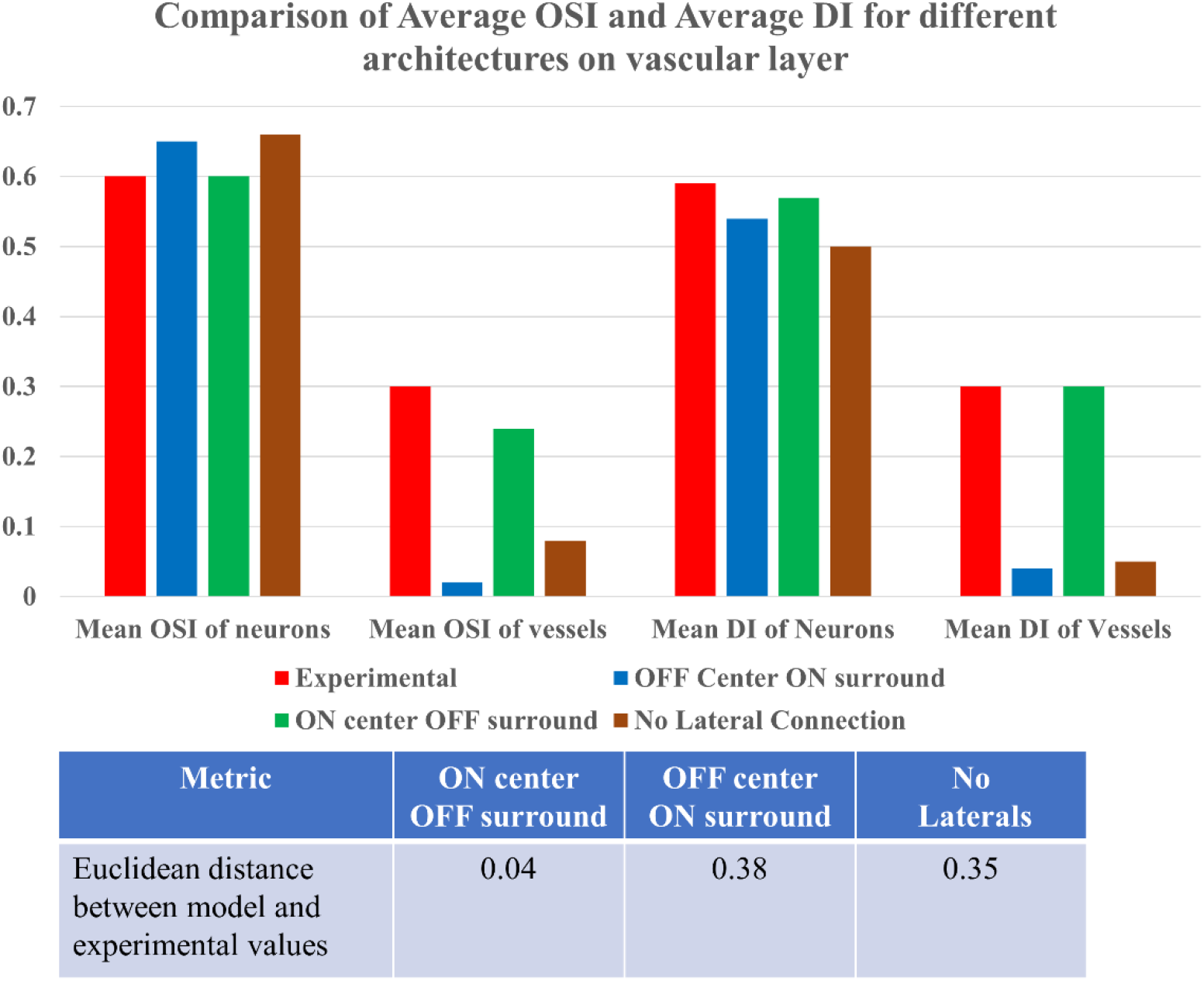
The comparison of model simulations with various architectures to the experimental values.

The OSI and DI of each vascular unit were estimated from the VLL. For each vessel, the perfusion field (~430 μm) surrounding it was identified in the NLL, and the average activity in that area was used to calculate the average neural OSI and DI. The orientation preference of the center vessels matched the orientation preference of the average activity of the neural units in their surrounding perfusion field (fig. 5). The distributions of the OSI in the neural and vascular units for all the three vascular topologies are compared in fig. 6. While all the three topologies resulted in a comparable value of OSI in NLL (~0.6), close to experimentally observed value [10], only the ON center OFF surround topology in VLL resulted in the OSI of vascular units close to the experimental observations (~0.3)[10] as seen in fig. 6c. In the right panel, the scatter plot shows that even though the blood vessels show tuned responses, their orientation selectivity is not correlated to the neural OSI (Pearson correlation coefficient = 0.15). The other topologies, No laterals (fig 6.a) and OFF center On surround (Fig 6.b), the vessels have weak to nil orientation selectivity.

**Figure 5:**
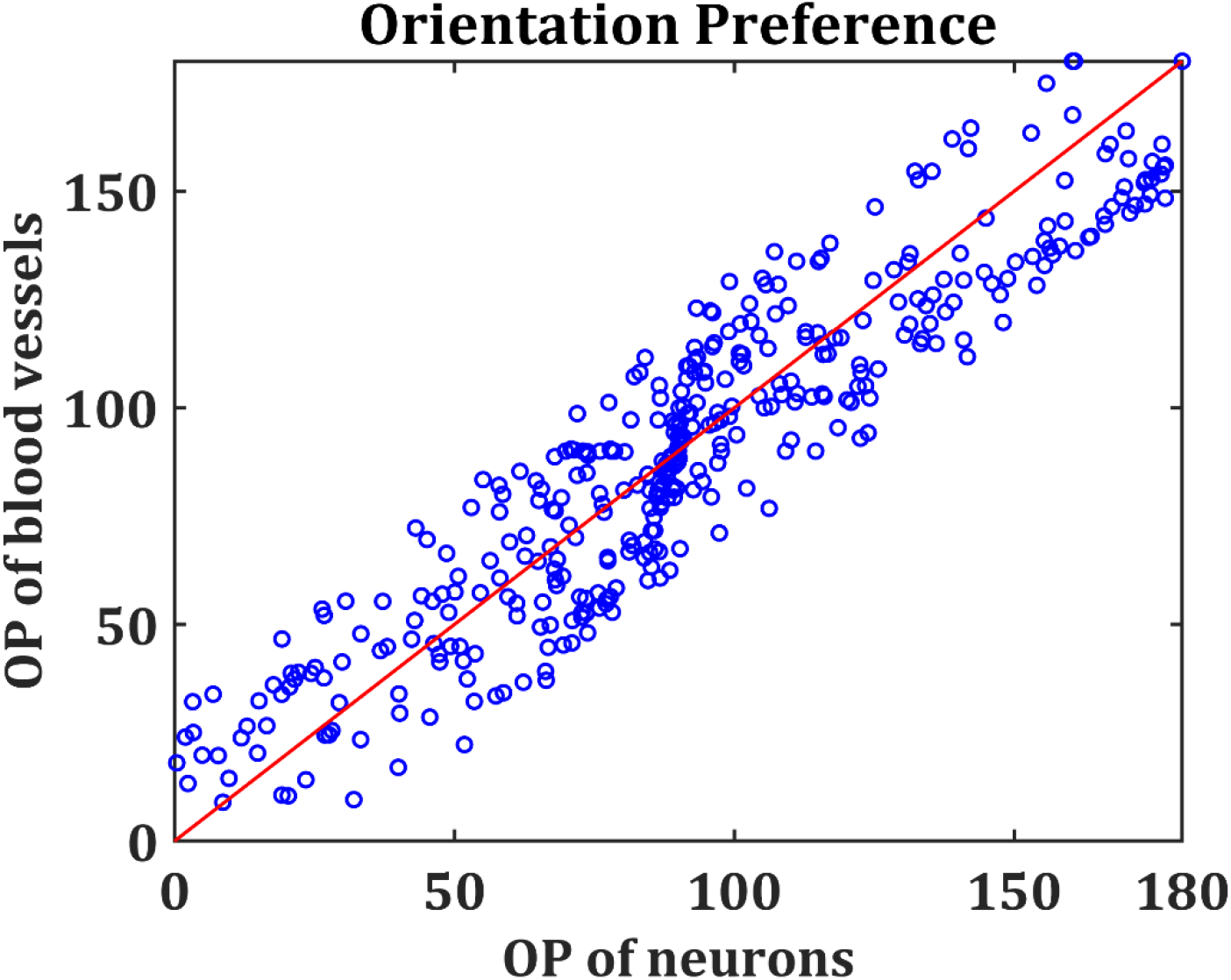
Orientation preference of vessels and the neural units around it (~430μm)

**Figure 6:**
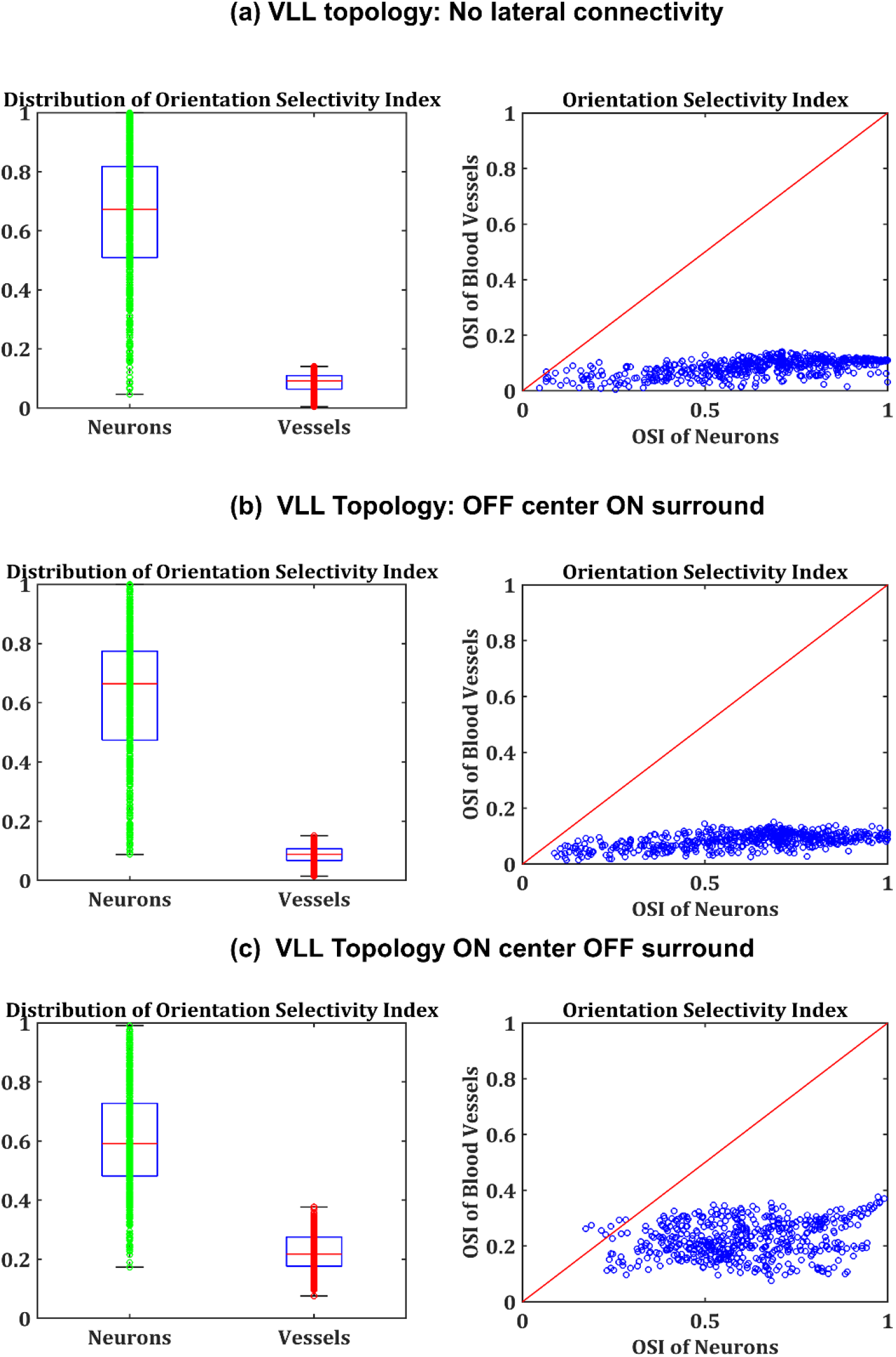
Left panel: The comparison of OSI in NLL and VLL by varying vascular toplogy. Right panel: The scatter plot of the OSI of NLL units and VLL units for the three VLL topologies (a) No laterals (b) OFF center ON surround (c) ON center OFF surround

Similarly, fig. 7 describes the DI of the VLL units with all three vascular topologies. Here also, it was observed that the ON center OFF surround topology (fig. 7c) resulted in a DI in NLL (~0.6) and in VLL (~0.3) closest to the value observed experimentally [10]. The other two topologies, No laterals and the OFF center ON surround resulted in a weak direction sensitivity in the vascular units. Similar to the case in OSI, the neural and vascular units showed no correlation (right panel of fig. 7c) with regard to the DI values (Pearson correlation coefficient = 0.14).

**Figure 7:**
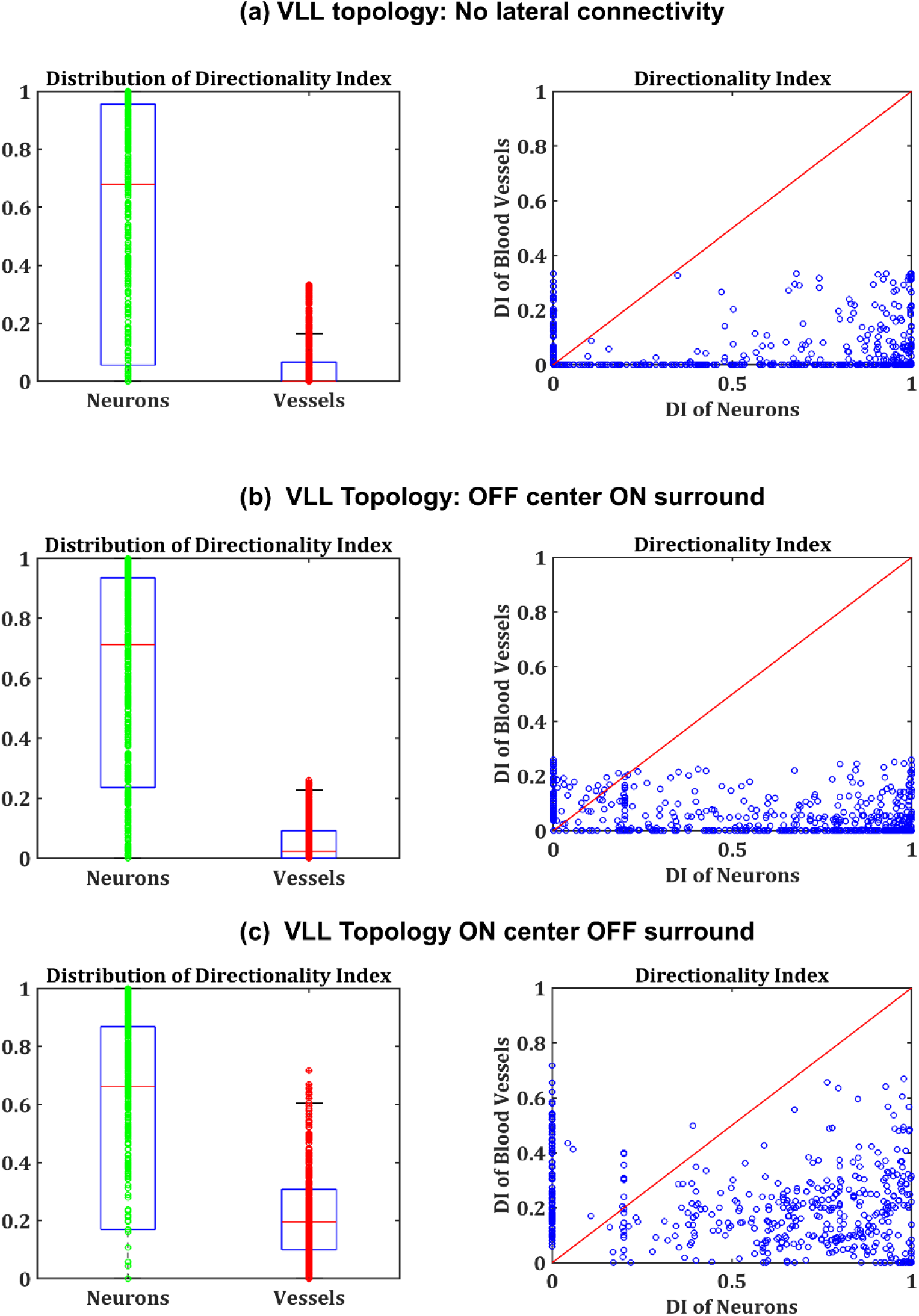
Left panel: The comparison of DI in NLL and VLL by varying vascular toplogy. Right panel: The scatter plot of the DI of NLL units and VLL units for the three VLL topologies (a) No laterals (b) OFF center ON surround (c) ON center OFF surround

The average neural activity was estimated by considering the perfusion field of the vessel to be ~430 μm in the figs. 6 and 7. This was inspired by the idea that occlusion of a single penetrating arteriole in the neocortex could result in tissue damage over an area around ~400 μm surrounding the arteriole [10,49]. But we also calculated the selectivity of neural responses by varying window size of averaging neural activity between 100 μm and 600μm following the experimental protocol adapted by O’Herron et al [10]. As seen in fig. 8, regardless of the window of neural activity considered, the selectivity of neural units was higher than that of the vascular units.

**Figure 8:**
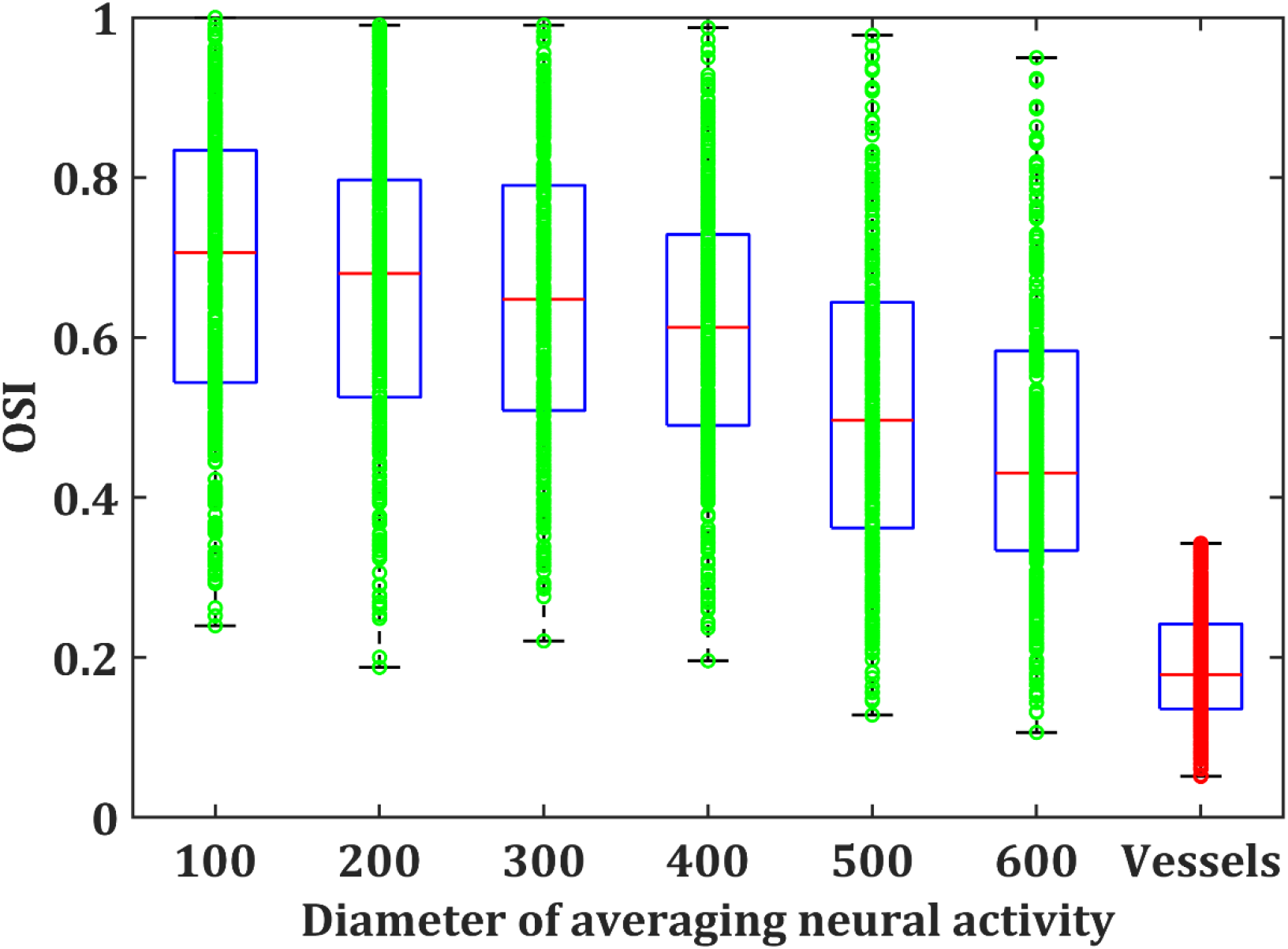
The OSI based on neural activity averaged over different diameters around the central vessel.

The activity of the NLL and VLL (ON center OFF Surround topology), in response to the drifting gratings in directions 45° and 135° is shown in fig.9. The left column shows the input stimulus, the middle column shows the NLL response to the input stimulus and the right column shows the response of VLL to the input stimulus. The vascular responses, although they follow the neural patterns, are clearly more diffuse and thus consistent with the experimental data shown by O’Herron et al.[10]

**Figure 9:**
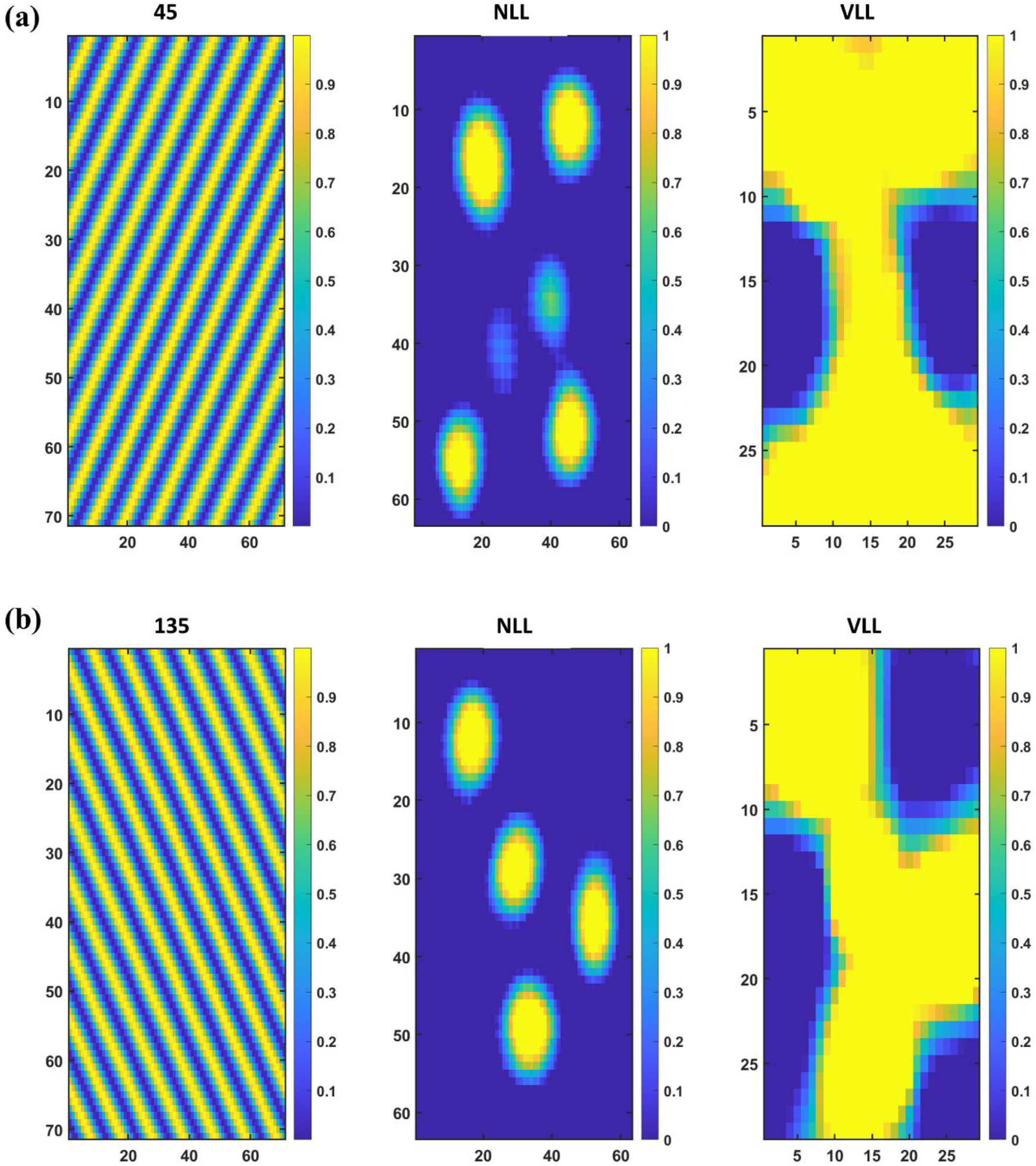
An example of activity in NLL and VLL in response to moving gratings with directions (a) 45° and (b)135°. The left column shows the input stimuli, middle column shows the activity of NLL and Right column shows the activity of VLL in response to the input stimulus.

### The influence of vascular feedback on the neural map formation

The vascular feedback to the neural network is critical to ensure the continuous energy supply to the neurons. The impact of vascular feedback on neural network characteristics has been explored by both experimental [52] and modeling studies [30]. The lateral connectivity and the tuned response in vessels define the vascular feedback to the neural layer in a totally different perspective. It implies that an alteration in the vascular feedback could change the way the neural layer encodes the input stimuli, thereby playing a critical role in the information processing.

The neural map characteristics are known to depend on the receptive fields of V1 neurons. The orientation map, especially in the visual cortex (V1), is observed to have a columnar organization or salt and pepper organization in different animals based on the ratio of the size of V1 to the size of the retina [53], or, more specifically, based on the size of the receptive field of V1 onto the retina. Keeping this in mind and considering that the neural responses are dependent on vascular feedback as well, we ask: “Would the perfusion field (PF) of the vessel affect the neural map formation?” To answer this question, we used bidirectional coupling between NLL and VLL, where internally both layers had ON-center, OFF-surround lateral connectivity. It was verified that the model reproduced the experimentally observed direction sensitivity values.

To observe the effect of the perfusion field of vessels on neural map formation, we varied the perfusion field from as small as 1×1 to as high as 40×40. The map was characterized as “columnar” or “salt and pepper” based on the attributes of the NLL’s response pattern to input stimuli. These attributes are defined as: (i) Number of Active Units (NAU) and (ii) Mean Area of Active Units (MAAU). When the perfusion field was small, the NAU was high, and the MAAU was small, indicating multiple activity blobs, each with a small area resembling a salt and pepper map (fig 10.b). As the perfusion field of the vessel increased, the NAU reduced while increasing the MAAU showing a transition to columnar map formation (fig. 10c). From fig. 10a, it is evident that neural networks with similar lateral connectivity architecture could encode the same input signal in a different way based on the perfusion field of the vascular network.

**Figure 10:**
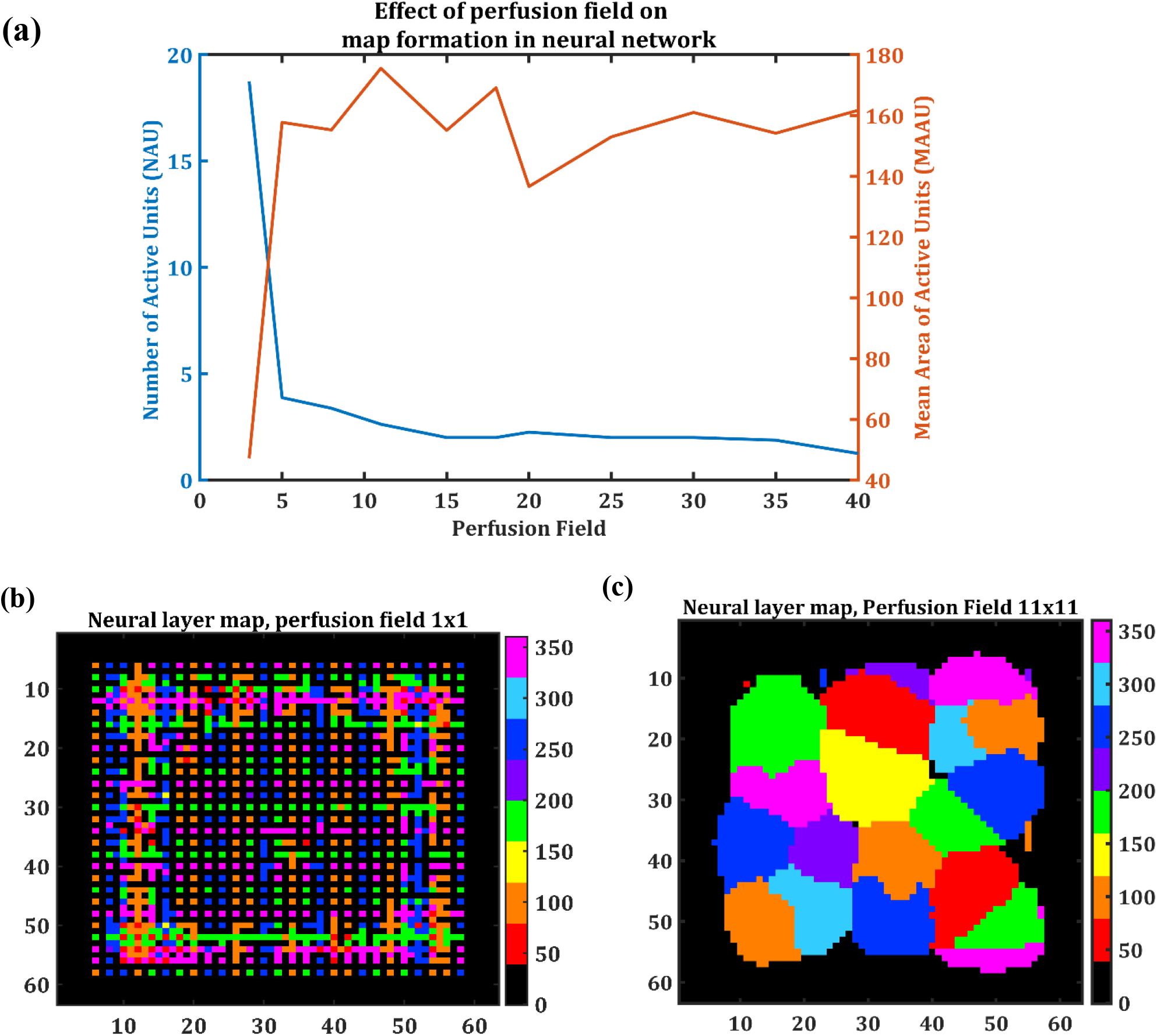
(a) The variation in the number of active neuron groups (blue) and the area of each neural activity group (red) in response to change in the perfusion field area. (b) The salt and pepper organization resulting from a small PF (1×1) and, (c) the columnar organization resulting from the larger PF (11×11). The black border in the surrounding region is due to the edge effect caused by taking limited size sheets to represent neural and vascular layers.

## Discussion

While information processing functions in the brain are primarily attributed to the neuronal network, the vascular network, which densely spans the entire brain area, is generally considered a passive irrigation system that ensures a timely supply of oxygen and nutrients to the neuronal network. It is generally assumed that the neural activity exerts a unidirectional influence on the vascular activity, rendering the vascular activity a faithful echo of the neural activity, an assumption that forms the basis of interpreting functional neuroimaging techniques [26,28,29]. In reality, the two networks (vascular and neural) are coupled in a bidirectional fashion, a fact that confronts the traditional view of leader-follower relationship between neural and vascular systems. There is a continuous flow of vasodilatory signals from the neurons to vessels and a continuous flow of energy substrates from vessels to the neural tissues. It is established by prior studies that any significant change in vascular feedback could alter the neural characteristics, especially during development [30,43,44,52]. The possibility of retrograde signaling from vessels to neurons was confirmed by Kim and colleagues[2] where the tone of the vessels influenced the firing rate of pyramidal neurons. Moore and Cao[1] predicted that the hemodynamics can potentially alter the gain of the cortical circuits. The computational model of Chander and Chakravarthy [43] also shows that hemodynamics can alter neural firing pattern by a retrograde influence. Hence the vascular network could play a significant role in modulating the information processing carried out by the neural network and therefore influence the long-term neural response to stimuli. In spite of this, the key aspects of an information processing network, such as tuned vascular responses were rarely sought out, until recently[10]. In their pioneering study, O’Herron and colleagues observed that the vessels in the cat visual cortex are capable of exhibiting tuned response to oriented moving gratings.

We modeled the development of a neurovascular coupling network by considering both the neural layer and the vascular layer as self-organizing networks with lateral connectivity (LISSOM) and introducing bidirectional coupling between them (fig.1). LISSOM has been used as a biologically plausible model to simulate the development of topographic visual maps [27]. The lateral connections with short-range excitatory projections and long-range inhibitory projections facilitate competitive learning and are ideal for modeling the retinotopic map formation seen in the primary visual cortex (V1). The lateral connections trained using asymmetric Hebbian learning [31] make the neural units naturally direction sensitive. There are advanced models of LISSOM like Gain-Controlled Adaptive LISSOM (GCAL) [54,55], which are more stable and robust. However, the threshold of the neural units in GCAL is adaptive and adapts depending on the average activity of the network. In NV-LIN, the threshold of the neural layer should depend on the vascular feedback – a feature that was earlier used in neurovascular models [44] and not the average neural activity in order to capture the effects of the vascular feedback. Hence, we used the LISSOM model and modified the equations in such a way that the vascular feedback is reflected in the output of the neural unit (eqn.1 and eqn.8).

The lateral connectivity of cortical neurons is known [27] to have an ON center OFF surround neighborhood connectivity. But the only hint that the tuned response in vessels gives is that there is a possibility of competitive learning among them, most likely brought about by inhibitory lateral connections [16]. To verify this, we varied the lateral architecture of vessels in the NV-LIN in three possible ways, (i) No lateral connectivity, (ii) ON center OFF surround, and (iii) OFF center ON surround. The resulting network was evaluated in terms of its OSI and DI and comparing it to the experimentally observed [10] OSI and DI in both neurons and vessels. The simulations led to the conclusion that the vessels are more likely to have lateral connectivity with an ON center OFF surround architecture similar to the neural network (fig.4). The ‘ON center’ signaling causing vasodilation in proximal vessels can be attributed to the endothelium-derived relaxing factors like nitric oxide [18]. The ‘OFF surround’ or vasoconstriction of distant vessels was observed by previous experimental studies[56]. The possible origin of these vasoconstrictions could be the activity of inhibitory interneurons [23,24] or astrocyte mediated pathways [57], which are capable of inducing vasodilation or vasoconstriction [25,26].

The role of vascular feedback in modulating the neural response is often overlooked while simulating the development of biological neural networks. The simulations from our NV-LIN model re-emphasize the relevance of vascular feedback in neural information processing. The OSI and DI exhibited by the neurons in our model matched the biologically observed values only when the vascular lateral connectivity architectures was ON center OFF surround. In addition, by varying the perfusion field of the vessel, our NV-LIN model demonstrated that the neurons respond differently to the same input stimulus. Note that there is considerable circumstantial evidence that the maturation of neural and vascular circuits is tightly coupled during early postnatal development [58,59]. Moreover, smooth muscle cells on blood vessels have molecules that serve as guidance cues for axons [60]. Quite remarkably, our simulations also suggest for the first time that the vasculature could also play a critical role in the formation of specific types of known functional architectures of neural maps in the cortex (see Fig. 10). Specifically, in our simulations, small perfusion areas result in a salt-and-pepper neural map (Fig. 10b). Presumably, small vascular perfusion areas may be sufficient for the metabolic needs in thin and small neocortical regions, e.g., mouse V1 (~750 um thick)[61] Our simulations further reveal that large perfusion areas produce highly clustered neural maps. We believe that more efficient functional hyperemia will occur when neural maps are highly clustered, as is seen in cat and monkey V1 and mouse barrel cortex which is ~ 1.75 mm thick.

Studies have shown that alterations in blood flow patterns and hypoperfusion are some of the early indications of several neurodegenerative diseases [63–65]. But the vasculature could play a critical role even at the onset of normal sensory experience in early postnatal development. It is possible that a salt-and-pepper orientation map organization might appear as a consequence of spatially-restricted blood flow in the vessels of V1, i.e., in non-rodent mammals such as cats, ferrets and monkeys; a spatially-restricted perfusion field could lead to a failure in the maturation of clustered and columnar cortical maps. This could be validated biologically by designing an experiment where the cortical map formation in early post-natal development is analyzed after varying the vascular perfusion area by using optogenetic techniques that can directly and selectively control the dilation in a subset of arteries[66]

Communication between microvascular networks is facilitated by mechanisms like hemodynamic coupling, diffusive and convective transport, and conducted responses [67]. Similar to brain regions, the metabolic demand patterns of other organs also exhibit considerable spatio-temporal variation [68]. Therefore, it is plausible that the nearby vessels should be able to sense the local energy demand of the metabolizing tissue and convey it to the larger proximal feeding and draining vessels. Dynamical interaction among the vessels which seeks to optimize microcirculation is thought to be the origin of a spatio-temporal phenomenon called vasomotion [69].

To conclude, our NV-LIN model of neuro-vascular map development confirms recent biological observations that sensory-evoked response in the vascular network is not merely a mirror of the tuned neural responses. The vascular network likely has an active ON-center OFF-surround lateral connectivity (equivalent to ON-center increases in blood flow vs. OFF-center decreases in blood flow) to display the tuned response observed biologically. Quite surprisingly, vascular characteristics such as the perfusion field of the vessels can impart significant influence on how functional architecture of the neural network develops. The cerebral microvessels, like neurons, are capable of having trainable connections and exhibiting competitive learning leading to tuned responses. Thus, vascular networks are likely much more than irrigation networks, and they might actually be critical modulators of neural map formation and information processing in the brain.

## Acknowledgments

We thank The Department of Biotechnology (DBT), Ministry of Science and Technology, Government of India (BIO/17-18/303/DBTX/SRIN), the Center for Complex Systems and Dynamics, IIT Madras, and NIH R01 MH111447 for partial funding of this project.

## Author Contributions

BSK did model designing, coding, analysis of the results, and manuscript preparation, VSC did model designing, analysis of the results, and manuscript preparation, P.O’H. and P.K were involved in analysis of the results, and manuscript preparation. All authors commented on and approved the final manuscript.

